# A complex copy number variant underlies differences in both colour plumage and cold adaptation in a dimorphic seabird

**DOI:** 10.1101/507384

**Authors:** Anna Tigano, Tone K. Reiertsen, James R. Walters, Vicki L. Friesen

**Affiliations:** Department of Biology, Queen’s University, Kingston, ON K7K 3N6, Canada; Norwegian Institute for Nature Research (NINA), FRAM - High North Research Centre for Climate and the Environment, 9296 Tromsø, Norway; Ecology and Evolutionary Biology, University of Kansas, Lawrence, KS 66046, USA; Present address: Molecular, Cellular and Biomedical Sciences, University of New Hampshire, Durham, NH 03824, USA

## Abstract

Colour morphs associated with different physiological adaptations offer unique opportunities to study the genomic basis of otherwise elusive adaptive traits. These complex balanced polymorphisms are often controlled by groups of tightly linked genes, and understanding how these ‘supergenes’ evolve and are maintained is an active area of research in evolutionary biology (Schwander *et al*. 2014, Thompson and Jiggins 2014). Within the Atlantic, the common murre (*Uria aalge*, a colonial seabird) displays a plumage colour dimorphism (‘bridled’ and ‘unbridled’) that seems to be associated with differences in thermal adaptation (Birkhead 1984; Reiertsen *et al*. 2012). The genes associated with bridling and how these genes affect thermal adaptation are unknown. Using whole genome resequencing, we investigate the genomic basis of differences in colouration and thermal adaptation between the two morphs, and how the association between the two traits is maintained despite random mating. We identify a 60 kb genomic region of high differentiation laying in the intergenic area amongst three candidate genes for colouration and thermal adaptation: *retinoic acid receptor beta* (*RARB*), *thyroid hormone receptor beta* (*THRB*), and *nuclear receptor subfamily 1 group D member 2* (*NR1D2* or *Rev-erbβ*). Differentiation is due to a complex copy number variant (CNV) that suppresses recombination locally. We show that this CNV acts as a ‘supergene’ and maintain association between regulatory elements likely affecting gene expression of one or more of the identified candidate genes. Our analyses also provide insights into the origin of the dimorphism: while copy number proliferation in the unbridled haplotype was potentially mediated by transposable elements (TEs), the bridled haplotype seems to have introgressed from the more cold-adapted sister species, the thick-billed murre (*U. lomvia*). Our results highlight the role of copy number variants in adaptation, especially when association among traits is maintained in the face of gene flow. They also shed light into the molecular mechanisms of adaptive thermogenesis in birds, which is poorly understood.

**Highlights:**

- Differences in plumage colour in Atlantic common murres are associated with different thermal adaptations
- A single region is highly differentiated between bridled and unbridled morphs
- A complex copy number variant in a non-coding region underlies the dimorphism
- Transposable elements and adaptive introgression from the thick-billed murre seem to explain the origin of the dimorphism

## RESULTS AND DISCUSSION

A central aim of evolutionary biology is to understand how diversity among and within species is generated and maintained. Colour is one of the most conspicuous form of phenotypic diversity. Colour polymorphisms, the co-occurrence of two or more colour morphs within a single interbreeding population, have been long studied to address central questions regarding sexual and natural selection. Colour polymorphisms can affect fitness directly, e.g. via sexual selection or predation avoidance, or indirectly, if the genes controlling the colour polymorphism are associated by either linkage disequilibrium (LD) or pleiotropy with genes underlying one or more traits that are the direct target of selection (McKinnon & Pierotti 2010). In the latter case, colour polymorphisms can be powerful markers for investigating otherwise cryptic phenotypic differences occurring within a species.

The common murre (*Uria aalge*) is monomorphic in the Pacific Ocean, but during the breeding season in the Atlantic Ocean it is dimorphic (Fig. 1): the bridled morph exhibits white plumage around the eye and down the auricular grove, whereas the more common unbridled morph has a completely black head. This dimorphism offers the opportunity to investigate the role of natural selection in maintaining phenotypic morphs despite random mating (Birkhead *et al*. 1980), because other factors known to play an important role in other species, such as aposematism, mimicry and sexual selection can be excluded in this study system (Kristensen *et al*. 2014). To confidently identify this dimorphism as adaptive, its effect on fitness and its genotype must be characterized (Barrett & Hoekstra 2011). Bridling seems to be associated with cold adaptation: the frequency of bridling correlates with sea surface temperature, with higher frequencies reported for colder temperatures at the breeding colonies (Birkhead 1984); and survival analyses indicate that survival increases with warmer temperatures in unbridled birds while it decreases in bridled birds (Reiertsen *et al*. 2012). Temporal trends in survival of the two morphs also indicate that the dimorphism is maintained by fluctuating, balancing selection (Reiertsen *et al*. 2012). Bridling behaves as a simple recessive Mendelian trait with the unbridled being dominant to the bridled morph (Jefferies & Parslow 1976). However, it is not known which genes underlie bridling and thermal adaptation nor it is clear how the association between the two traits evolved and is maintained.

**Figure 1.**
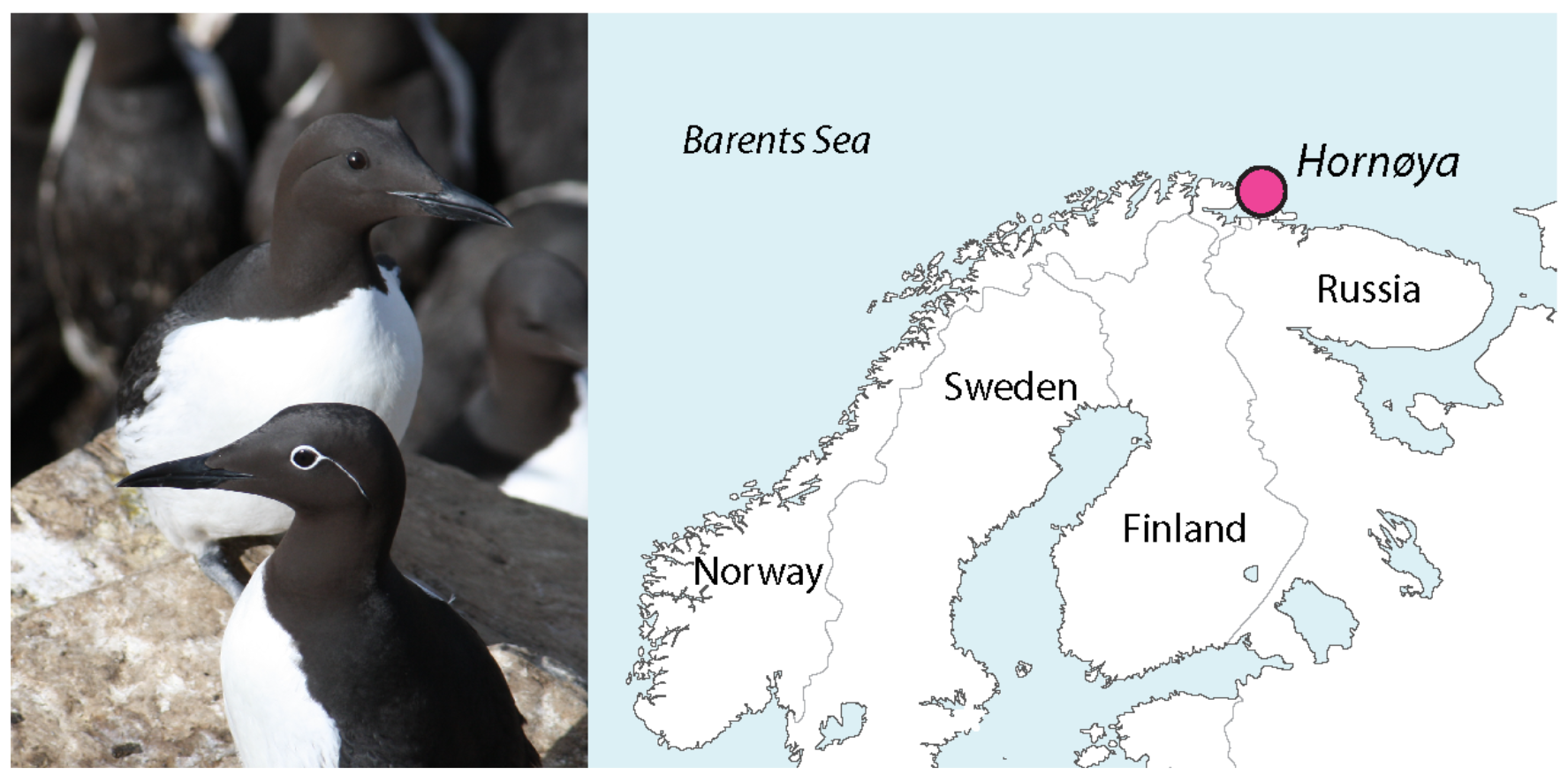
Picture showing an unbridled (above) and a bridled (below) common murre, and the geographic location of sampling.

### A single genomic region of high differentiation between morphs

Unless the two traits are controlled by a single locus or strong selection alone is acting on both traits, gene flow is expected to disrupt the association between bridling and cold adaptation if recombination is not reduced or suppressed (Tigano & Friesen 2016). To investigate the genomic basis and apparent association of bridling and cold adaptation, we sequenced the whole genome of 10 bridled and 10 unbridled common murres from the colony at Hornøya, Norway, where the frequency of bridling is ~30% (Fig. 1). Using Illumina 150 bp paired-end reads, we generated ~8-fold genome coverage for each sample, and mapped the reads to a consensus reference genome based on the thick-billed murre (*Uria lomvia*, sister species of the common murre) reference genome (Tigano *et al*. 2018). We calculated genome-wide differentiation using 50 kb moving windows using FST and identified two windows on scaffold 72 that had the most extreme FST values (0.55 and 0.51, based on common variants; MAF>0.1) and that were in the 99.9 quantile of the genome-wide FST distribution (average FST=0.09; Fig. 2A). To test whether the genotypes of individuals were consistent with predictions based on phenotype (Jefferies & Parslow 1976) and to test for deviations from a model of neutral evolution, we calculated FST, nucleotide diversity (π) and Tajima’s D using 10 kb moving windows within 1 Mb of the area of differentiation. FST showed high values along a 60 kb span (Fig. 2B). In the area corresponding to the FST peak, π was much higher in unbridled birds compared both to bridled birds and to the surrounding genomic area (Fig. 2B). Similarly, Tajima’s D estimates were positive in unbridled birds, consistent with balancing selection, and negative in bridled birds, consistent with a selective sweep (Fig. 2B), and provided evidence that both haplotypes differed from the surrounding background genome independently from one another. Although estimates of π and Tajima’s D are consistent with the unbridled morph being dominant over the bridled morph, π in unbridled birds exceeded expectations based on unbridled birds being heterozygous only (see below).

**Figure 2.**
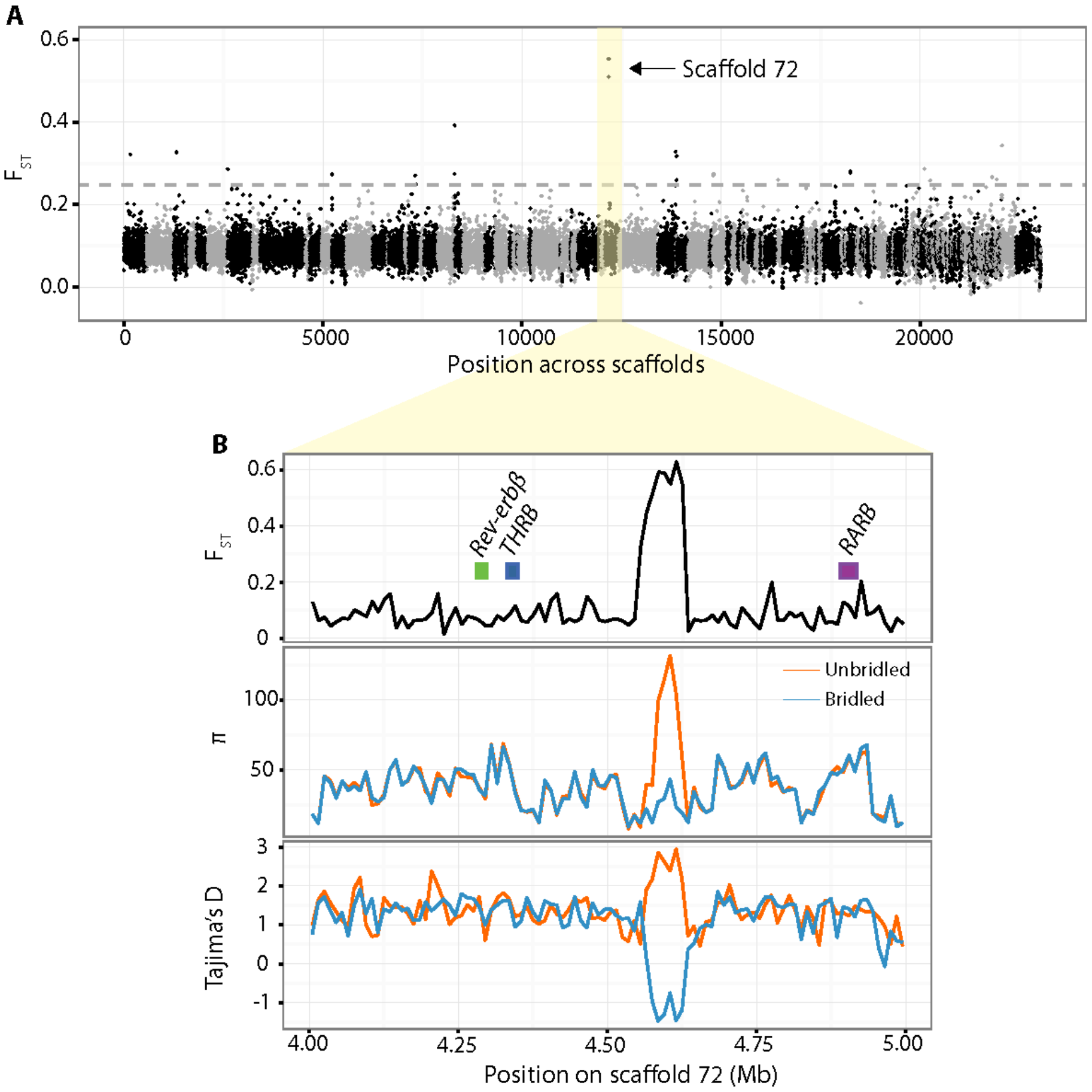
Patterns of differentiation and genetic diversity. A) Manhattan plot showing genome-wide differentiation using 50 kb moving windows. Dashed line indicates 99.9^th^ percentile of the empirical distribution. In yellow are highlighted the two windows on scaffold 72 showing the most extreme FST values. B) Plots showing FST with position of closest genes, nucleotide diversity (π) and Tajima’s D in 10 kb moving windows within 1 Mb of the area of differentiation.

### Suppressed recombination in the area of differentiation

Analyses of differentiation at the single nucleotide polymorphism (SNP) level identified three distinct blocks of high differentiation (Fig. 3A). To test whether recombination among these blocks was suppressed, we calculated LD among SNPs within 200 kb of the FST peak. Blocks of high differentiation matched blocks of high LD (Fig. 3A; Fig. S2): when compared to the surrounding genomic area, LD in the FST peak area was high not only among SNPs within a genomic block, but also among blocks, suggesting that recombination within and among blocks is suppressed despite random mating between the morphs, and that the whole area of differentiation is inherited as one genomic block (Fig. 3B, Fig. S2). We observed the greatest difference in LD between peak vs. no peak regions in unbridled birds (Wilcoxon signed-rank test: p-value < 0.0001; Fig. 3B). Assuming high synteny among avian genomes (Ellegren 2010), we could exclude that suppressed recombination was due to proximity to a centromere (Tigano & Friesen 2016) because the area of differentiation mapped 15 Mbp away from the centromere of chromosome 2 in the chicken genome. Therefore, we hypothesized that recombination was suppressed due to a structural variant, such as an inversion. Read depth was inflated by 3- to 11-fold in unbridled birds in the area corresponding to the FST peak (Fig. 3C), pointing to the presence of a copy number variant (CNV) in unbridled birds when compared to bridled birds and the reference genome, rather than an inversion. Over-hanging orphan reads at the edges of the area of differentiation, and additional analyses of read pair orientation and read depth indicated that the CNV was a complex structural variant (see Supplementary Material). A large deletion (15 kb) detected between LD blocks 1 and 2 (Fig. 3A, 3C), and the extent of the differences in nucleotide diversity (Fig. 2B) and read depth (Fig. 3C) between morphs could not be explained by either heterozygosity alone, or a simple duplication event, and support presence of multiple copies. As recombination efficiency decreases with INDEL size (Kung et al. 2013), the extent of the CNV (up to ~600 kb) appears to be the sole factor affecting recombination between the two haplotypes. However, since the breakpoints of the CNV variants could not be fully characterized, we cannot exclude additional structural variants, such as inversions, or other genomic mechanisms contributing to recombination suppression between the two haplotypes (Tigano & Friesen 2016).

**Figure 3.**
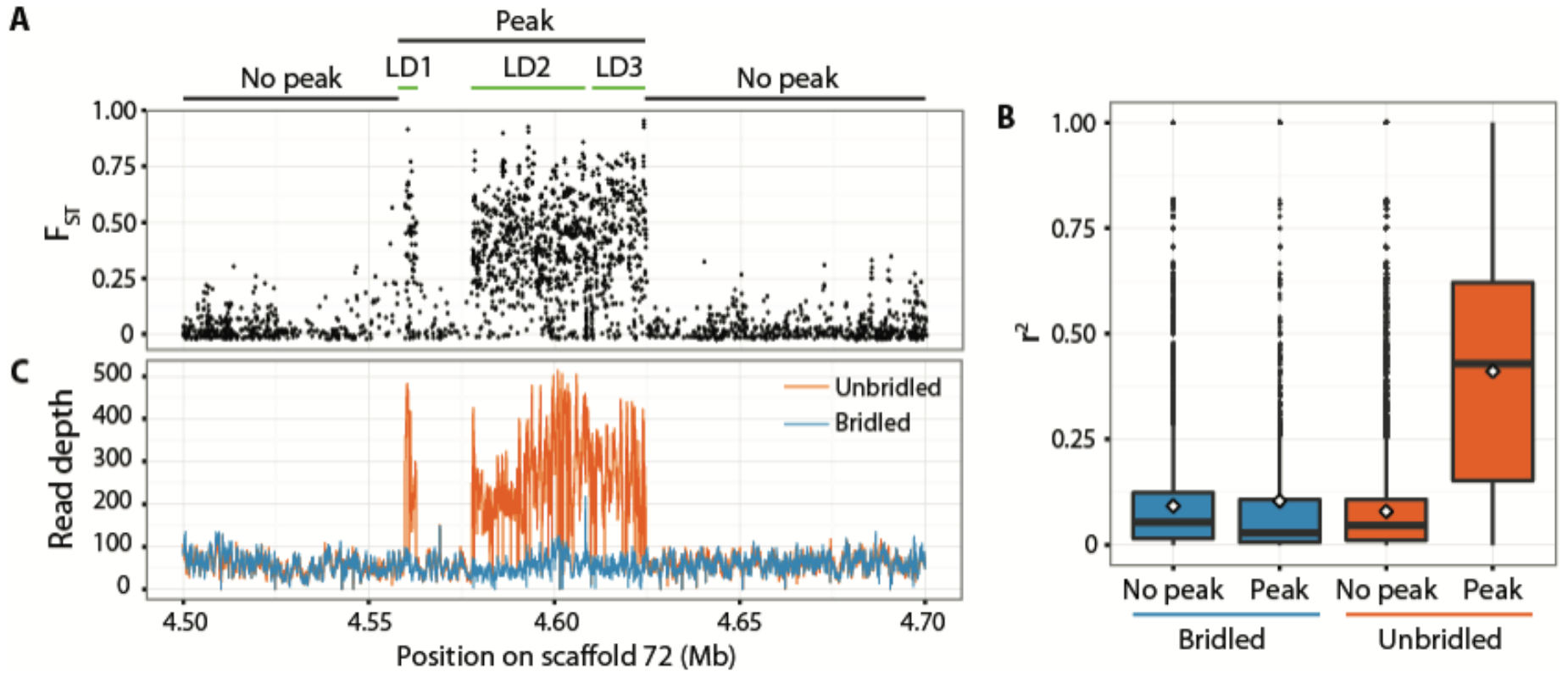
Fine-scale differentiation, patterns of LD and read depth in the area of differentiation and surrounding regions. A) Each dot in the plot represents a single SNP FST estimate. Genomic blocks of differentiation match blocks of LD (LD1-3; Fig. S2). B) Levels of LD for each peak and no peak area in the 200 kb surrounding the area of differentiation for each of the two morphs. Black lines indicate the median, white diamonds indicate the average. All comparisons were significantly different (Wilcoxon ranked-sum test: p-value<0.001). C) Read depth for the same 200 kb region. Positions of inflated coverage in unbridled birds match closely the positions of high differentiation in A).

### Candidate genes for differences in colour and/or thermal adaptation

The 60 kb area of differentiation lacked annotated genes (Fig. 2B). We propose that differentiation in intergenic regulatory regions, likely involving copy number variants and perhaps sequence divergence, affect expression of neighbouring genes (Spielmann and Klopocki 2013), thus explaining phenotypic differences between the two morphs. This hypothesis is supported by the presence in this area of several non-coding regions that are highly conserved across birds and mammals (Fig. S1A) potentially associated with gene regulatory elements (Hardison 2000). Additionally, enhancers and active regulatory elements were identified in the corresponding area in the human genome, particularly for epidermal keratinocytes and skeletal muscle myoblasts (Fig. S1B; ENCODE Project Consortium 2012). In birds these cell types are respectively the precursors of feathers and skeletal muscle, the main avian thermogenic organ (Newman *et al*. 2013). Multiple copies of regulatory elements in the intergenic area presumably affect gene expression of neighbouring genes and connect genotypic differences to phenotypic differences between bridled and unbridled common murres.

We identified three genes whose changes in expression could explain bridling and/or cold adaptation in bridled murres (Fig. 2B). The first was *RARB*, a retinoic acid receptor, located 280 kb downstream of the FST peak. *RARB* is up-regulated during feather development (Ng *et al*. 2015), and its ligand retinoic acid is a potent inhibitor of tyrosinase (Orlow *et al*. 1990), a key enzyme in melanin synthesis, which identifies *RARB* as a candidate gene for bridling. *RARB* is a highly pleiotropic gene and is involved in a multitude of biological processes, including thermogenesis. The first two annotated genes upstream of the FST peak, 210 and 250 kb respectively, were *THRB*, a thyroid hormone receptor, and *Rev-erbβ*, a transcriptional repressor, both of which also are involved in thermogenesis (Lowell & Spiegelman 2000; Gerhart-Hines *et al*. 2013). In birds, thyroid hormones, in particular triiodothyronine (T3), regulate basal temperature and resting metabolic rate (Elliott *et al*. 2013), and high circulating levels of T3 have been associated with thermoregulation in response to cold challenges (Vézina *et al*. 2015). *Rev-erbα* and *Rev-erbβ*, act in concert as important regulators of the circadian clock and energetic homeostasis (Bugge *et al*. 2012; Cho *et al*. 2012; Solt *et al*. 2012). Knock-out experiments have shown that Rev-erbα-null mice have better survival than normal mice at cold temperatures (Gerhart-Hines *et al*. 2013).

Adaptive thermogenesis, the production of heat in response to cold challenges, is a complex biological process (Lowell & Spiegelman 2000), and while it has been studied extensively in mammals, the molecular mechanisms underpinning thermoregulation and thermogenesis in birds are not well understood. In mammals, *RARB, THRB* and *Rev-erbβ* regulate the expression of UCP-1, a mitochondrial uncoupling protein that enables the generation of heat in brown adipose tissue (Lowell & Spiegelman 2000). However, birds and other reptiles lack both brown adipose tissue and the gene encoding UCP-1 (Newman *et al*. 2013). The avian homologue of UCP-1 (avUCP) may compensate for the loss of UCP-1, although whether avUCP has the same thermogenic role or is mainly involved in protection from oxidative stress is unclear (Talbot *et al*. 2004). Nonetheless, *Rev-erbα* was shown to be differentially expressed in *Anolis* lizards adapted to different thermal environments (Akashi *et al*. 2016), suggesting that the genes associated with thermogenesis in mammals could have similar functions in birds and non-avian reptiles.

Our analyses strongly suggest that the association between colouration and adaptive thermogenesis in bridled and unbridled common murres is maintained via linkage between regulatory elements, rather than between coding candidate genes, and that those regulatory elements affect two different traits by controlling gene expression of one or more candidate genes in two different target tissues. *RARB* could act pleiotropically on the two traits either directly, through altered levels of retinoic acid in both target tissues (biological pleiotropy), or indirectly, if retinoic acid affected the expression of *THRB* and/or *Rev-erbα* (mediated pleiotropy). Alternatively, colouration could be controlled by *RARB*, and adaptive thermogenesis could be controlled by *THRB* and/or *Rev-erbα*, independently from one another. Further research is necessary to disentangle the role of each gene in thermogenesis.

Dominance of the unbridled morph, as established through hand-rearing experiments (Jefferies and Parslow 1976), combined with our results suggest that bridling could be due to a threshold effect associated with the expression of *RARB*. Whether cold tolerance associated with bridling is also a recessive trait, or if bridled/unbridled heterozygotes express an intermediate thermal tolerance phenotype remains to be investigated. The variant with high number of copies was found in five unbridled individuals, which is consistent with the frequency of bridling at this colony (half of the unbridled individuals are expected to be heterozygous given that frequency of bridling is 30% at this colony), and supported a significant association between the phenotype and the genotype based on Hardy-Weinberg expectations (Fisher’s exact test, p < 0.05). However, the rest of the unbridled birds (n=5) did not show an intermediate, heterozygous genotype as expected, presumably due to the inherent difficulties of genotyping complex structural variants in heterozygous individuals (Zhou *et al*. 2011). To characterize the CNV, we *de novo* assembled the genome of an unbridled common murre with high number of copies in the area of differentiation (Supplementary Material). Despite high overall contiguity (N50=12 Mb, Table S2), the resulting assembly was discontinuous in correspondence of both CNV breakpoints, and the area of high differentiation comprised short contigs (< 500 bp) with sequencing depth elevated 10-fold. Although these results prevented us from characterizing the CNV any further, they confirm the position of the CNV and the high number of copies that characterize the unbridled haplotype.

### Evolutionary origin of the dimorphism

Transposable elements (TEs) are known to promote chromosomal rearrangements, including duplications and copy proliferation, via homologous recombination or alternative transposition (Gray 2000). We tested whether an enrichment in TEs in the area of differentiation could explain the copy proliferation in the unbridled haplotype. TEs in this region were not more abundant than the genome average, but a 10 kb stretch at position 72:4,600,000-4,610,000 (i.e. in the area of differentiation) contained a significant, ~3-fold enrichment of TEs (18.3% vs. 6.5%, Chi-squared p < 0.001), where 80% of TEs sequences were LINE-class TEs. These results would suggest that bridling is the ancestral phenotype, from which the unbridled morph evolved.

However, because bridling is absent from the Pacific, the dimorphism probably evolved after divergence of the Atlantic/Pacific populations, 56,000-226,000 years ago (Morris-Pocock *et al*. 2008) with bridling rather being the derived phenotype. Nonetheless, a single mutation wiping away all but one copy of a large genomic area is highly unlikely to explain the evolution of bridling from the ancestral unbridled morph. Alternatively, the bridling haplotype could have been present in the Atlantic at low frequency or have introgressed from a different population or species. Indeed, adaptive introgression from the thick-billed murre is supported by several observations: a significant reduction in nucleotide diversity (t-test, p-value < e^-10^) and Tajima’s D values in the bridled haplotype (Fig. 2B) compared to genome average, and the presence of a single copy of the differentiated region in the thick-billed murre reference genome. Thick-billed murres are exposed to significantly lower temperatures than common murres throughout the year, and this difference has been implicated in divergence of the two sister species (McFarlane Tranquilla *et al*. 2015). Additionally, thick-billed and common murres are known to hybridize in both the Atlantic and Pacific Oceans (Taylor et al. 2012; Colston-Nepali 2017). Repeated backcrossing to common murres following hybridization may have purged thick-billed murre DNA with the exception of the bridled haplotype, due to a combination of its fitness effects and its intrinsic resistance to recombination. Additional data from both murre species are necessary to reconstruct the history of gene exchange between them, and further investigate the hypothesis that adaptive introgression lead to the evolution of the adaptive dimorphism in Atlantic common murres. Nonetheless, these results combined suggest that TEs might have had a role in the copy number proliferation and thus in the adaptive differentiation between common and thick-billed murres and/or between bridled and unbridled common murres.

### Conclusions

Our results indicate that a CNV can have an important role in local adaptation and can in fact act as a ‘supergene’: this large structural variant suppresses recombination between the two haplotypes, and thus maintains a complex balanced dimorphism despite random mating between the two morphs. The scarce knowledge on adaptive thermogenesis in birds hinders our understanding of how birds adapt to their thermal environment, and hence limits our ability to predict their adaptive potential to climate change (Angilletta 2009). In this study, we have identified not only a new candidate gene for plumage colouration (*RARB*), but also a set of candidate genes for thermal adaptation, which represent an important step towards the understanding of the genomic and physiological basis of adaptive thermogenesis in birds, and potentially other animals as well. Physiological studies stemming from these findings will clarify the role of each of the three candidate genes in adaptive thermogenesis and will advance our understanding of how species respond and adapt to thermal changes in their environment.

## Accession Numbers

Whole genome resequencing data for the 20 common murre individuals and the genome assembly of the unbridled common murre will be uploaded on NCBI.

## Supplemental Information

Supplemental methods and figures can be found in the Supplemental Information file.

## Authors contributions

AT conceived the idea for the study with input from TKR and VLF. AT and TKR collected samples. AT supervised sequencing and performed all analyses with guidance from JRW and VLF. AT wrote the manuscript. All authors contributed to the interpretation of results and provided critical feedback on the research, analysis and final manuscript.

## Acknowledgements

The authors thank McGill University and Genome Quebec Innovation Centre for whole genome library preparation and sequencing, and the Centre for Advanced Computing at Queen’s University for assistance with the computing cluster. Leonardo Campagna, Rob Colautti, Allison Shultz and Virginia Walker provided suggestions on analyses and helped with interpretation of results. Greg Robertson facilitated funding for the sequencing and assembly of the common murre genome. This project was funded by Natural Sciences and Engineering Research Council of Canada (Discovery Grant to VLF), Queen’s Graduate Research Awards (AT), American Ornithologists’ Union Research Grant (AT), Frank M. Chapman Memorial Fund (AT), Norwegian Institute for Nature Research (TKR) and Environment and Climate Change Canada (VLF and AT).

