## Supplemental Information for "A complex copy number variant underlies differences in both colour plumage and cold adaptation in a dimorphic seabird"

### Supplementary Material

#### Whole genome resequencing

We collected blood samples from the brachial vein of 10 bridled and 10 unbridled common murres (*Uria aalge*) from Hornøya, Norway, during summer 2014. After genetic determination of sex, we selected five females and five males for each morph to ensure that an equal number of individuals per sex were sequence for each morph. We extracted DNA from blood using a standard protease K/phenol-chloroform protocol (Sambrook *et al.* 1989), and purified DNA with ethanol precipitation followed by resuspension in DNAse-free water. Paired-end libraries were prepared for each individual using Illumina TruSeq DNA sample Prep Kit v2 with an average insert size of 270-420. Paired-end sequencing (150 bp) of 20 individual libraries on an Illumina HiSeq2500 generated a total of 185 Gb of data. Library preparation and sequencing were performed at the Genome Quebec Innovation Center (McGill University, Montreal). Average raw read sequencing coverage was 7.8X (6.2-12.0X) assuming a genome size of 1.18 Gb (Tigano *et al.* 2018).

#### Consensus reference genome

To align common murre whole genome resequencing reads we used a common murre consensus reference genome built from the reference sequence of the thick-billed murre (*Uria lomvia*), the sister species of the common murre (Tigano *et al.* 2018). We first mapped raw common murre reads to the thick-billed murre reference genome with BWA (Li & Durbin 2009) using the BWA-MEM algorithm, and included information on sequencing lane, sequencing run and individual to the alignment files. We marked and removed PCR duplicates with MarkDuplicates in PICARD TOOLS twice, once before merging alignment files for each individual and once after.

All following analyses were performed in GATK (McKenna *et al.* 2010) according to GATK Best Practices recommendations (DePristo *et al.* 2011; Van der Auwera *et al.* 2014) to build a consensus reference. First, we executed local realignment in regions around insertions and/or deletions (indels) to correct for the misalignment of bases using RealignerTargetCreator and IndelRealigner. We called variants with HaplotypeCaller and GenotypeGVCFs and filtered for quality, separately for the SNPs call set and the indels call set. We applied hard filtering as recommended, plus a minor allele frequency (MAF) filter of 0.2 to ensure that the reference sequence was replaced with variants that were common in common murres. Finally, we built the common murre consensus reference with FastaAlternateReferenceMaker by replacing the thick-billed murre sequence with one of the common murre variants for each polymorphic site.

**Genomic analyses of bridled and unbridled common murres**

*Variant calling and genotyping*

We aligned common murre raw reads to the common murre consensus reference, and processed mapped reads as described above (PCR duplicates removal followed by local realignments around indels). To call variants in common murre individuals, however, we adopted the approach implemented in ANGSD (Korneliussen *et al.* 2014), which is based on an algorithm specifically developed to accommodate the genotype calling uncertainty associated with low coverage data. Instead of calling genotypes directly, ANGSD calculates the Site Frequency Spectrum (SFS), i.e. the distribution of sample allele frequencies, based on individual genotype likelihoods (GL) for each site. Population genetic parameters can be estimated directly from the SFS and with higher accuracy because genotype likelihoods are taken into account. We proceeded in an iterative fashion. First, we calculated site frequency likelihoods (-doSaf 1) jointly for all samples based on GLs calculated with the Samtools model (-GL 1). We retained only polymorphic sites ($p<10^{-6}$) with minimum mapping quality of 30, minimum base quality scores of 20 and MAF > 0.10. We recalculated site frequency likelihoods for each morph separately for the polymorphic sites that passed the above filters and that were genotyped in a minimum of five individuals. Our dataset included 12,373,865 SNPs. We then ran realSFS in ANGSD to obtain a maximum likelihood estimate of the folded SFS using the Expectation Maximization (EM) algorithm for each morph separately and jointly for the two morphs (2dSFS).

*Genetic diversity and differentiation*

We calculated genome-wide $F_{ST}$ based on the 2dSFS using 50 kb non-overlapping moving windows across each scaffold to estimate genome-wide differentiation between the two morphs and to identify areas of high differentiation. We observed noisy window-based $F_{ST}$ estimates for scaffolds smaller that 50 kb, which we therefore excluded. We obtained a total of 23,028 high quality 50 kb windows for downstream analyses. Per-window $F_{ST}$ ranged from -0.0382 to 0.5534 and averaged 0.0891 across windows. Global $F_{ST}$ across all SNPs showed a similar value (0.0887). The relatively high global $F_{ST}$ value between bridled and unbridled common murres is likely due to the exclusion of rare variants from the dataset (MAF>0.10). We predicted a homogenous genomic background due to random mating between the two morphs (Kristensen *et al.* 2014) and one or few areas of high differentiation underlying the genes controlling the dimorphism. According to predictions, we identified two windows on scaffold 72 that showed extreme differentiation compared to the rest of the genome. To increase definition, we recalculated $F_{ST}$ in smaller moving windows of 10 kb for the one million bases surrounding the two $F_{ST}$ peaks (scaffold72:4,000,000-5,000,000). Considering that bridling is a Mendelian recessive trait, and that the frequency of bridling at Hornøya is ~30%, 50% of individuals should be heterozygous at that allele. Therefore, indices of genetic diversity should be higher across unbridled birds because at least some individuals should be heterozygous for the bridling allele. We calculated nucleotide diversity ($\pi$) as a measure of genetic diversity, and Tajima's D to test whether high differentiation in scaffold 72 showed a signature of selection. Tajima's D tests for deviations from a neutral model of evolution, including selection and demographic processes. The symmetrical deviations from neutrality in bridled vs. unbridled common murres in the area of high differentiation, with positive values in unbridled and negative values in bridled individuals, combined with a neutral distribution of Tajima's D in the surrounding area, are supportive of selection rather than demographic processes. However, background Tajima's D was slightly positive. Although positive values of Tajima's are indicative of loss of rare alleles and could indicate population contraction, these values could be inflated by removal of rare variants through filtering alleles with MAF<0.10.

*Linkage disequilibrium*

We phased 2,350 SNPs across 200 kb spanning the area of differentiation (scaffold 72:4,500,000-4,700,000) for each morph separately with beagle v. 4.1 (Browning & Browning 2007). We used Haploview (Barrett *et al.* 2005) to calculate pairwise linkage disequilibrium (LD) from phased genotypes for all samples together, and for each morph separately. The area of differentiation showed high LD distributed across three main LD blocks separated by areas of low LD. Using the abrupt changes in LD at the edges of the $F_{ST}$ peaks, we identified two main breakpoints at position 4,559,870 and 4,624,381, respectively. Internal areas of low LD were situated at positions 4,562,777-4,577,904 and 4,608,219-4,608,525.

Analyses of LD on each morph separately showed that LD, measured as $r^2$, was higher in the area of differentiation than in the surrounding area in both morphs, although the difference was small in bridled birds and large in unbridled birds (Table S1, Fig. S2). Differences in LD between bridled and unbridled common murres for the whole area, and between peak and surrounding areas in each morph, were all significant (Wilcoxon rank sum test: p-value < $e^{-10}$).

*Structural variant characterization*

High LD indicates a reduction or suppression of recombination. Blocks of LD have often been attributed to chromosomal inversions that suppress recombination between heterokaryotypes (Tigano & Friesen 2016). Thus, if an inversion were present, read depth should drop around the breakpoints of the area of differentiation. We plotted read depth for each of the two morphs and found that read depth was highly inflated in unbridled common murres in the region corresponding to high $F_{ST}$ SNPs, rather than dropping around the breakpoints, suggesting the presence of copy-number variants (CNV) due to duplication, rather than an inversion (but see below). To identify homozygote and heterozygote unbridled birds based on fold-differences in copy number, we plotted read-depth for each individual separately to identify homozygous and heterozygous unbridled common murres, and found that only 5 unbridled birds showed high coverage in that area. This is consistent with the frequency of bridling at the colony where the birds were sampled, but we failed to detect intermediate heterozygous genotypes.

Although high-throughput sequencing has contributed greatly to the characterization of structural variants (SV), such as inversions, deletions, insertions, CNV and translocations, in model and non-model species, analysis of SV using short reads still presents several limitations (Tattini *et al.* 2015). To increase our power in genotyping the SV associated with bridling, we used an integrative approach that combines different methods. We visually inspected alignment files with the Integrative Genome Viewer (IGV; Robinson *et al.* 2011; Thorvaldsdóttir *et al.* 2013) to identify read pairs with anomalous insert size and orientation. We then analysed sequencing data using four programs that are each based on different approaches: CNVnator (Abyzov *et al.* 2011) relies on read-depth to genotype CNVs specifically; Breakdancer (Chen *et al.* 2009) is based on a read-pair algorithm and is able to detect a variety of SV; Pindel (Ye *et al.* 2009) and Delly (Rausch *et al.* 2012) combine read-pair and split-reads methods to increase sensitivity and specificity. Breakdancer and Delly did not detect any SV in the area of differentiation. However, analyses of read pair insert size and orientation in IGV indicated that many read pairs had reversed orientation and were longer than average insert, with each read of the pair mapping at opposite ends of a 15 kb sequence, respectively, between LD blocks 1 and 2. Reversed orientation of read pairs indicates duplication, while larger than average insert size indicates deletion. However, read depth in that 15 kb area was similar to the genome average, indicating that the gap of inflated coverage rather represented lack of duplication in that area. These results were corroborated by CNVnator, which showed evidence of a duplication spanning 47 kb across LD blocks 2 and 3 ($p$-value < 0.0001), and Pindel, which indicated a deletion between LD blocks 1 and 2. Pindel identified a total of 69 SV, proving to be the most sensitive approach. Of these, only 4 were found in bridled common murres, and only 2 in unbridled common murres showing normal coverage in the area of differentiation, while the rest were exclusively found in unbridled common murres displaying inflated coverage at the $F_{ST}$ peak. Most SVs (58 out of 69) were small (<10 bp). In addition to the 15 kb deletion discussed above, Pindel identified another medium size deletion (767 bp) at positions 4,618,387- 4,619,154. All SVs genotyped in Pindel were confirmed by visual inspection in IGV. Note, however, that because the copy number variant in unbridled birds represents presumably a large insertion, the precise characterization of the variant is hindered by a reference genome that does not include this insertion (Kidd *et al.* 2010). Therefore, presence of an inversion that would explain the suppression of recombination in the area of differentiation cannot be excluded at this point.

***Common murre reference genome***

To better characterize the CNV in unbridled common murres, we *de novo* assembled the genome of one of the original unbridled individuals that showed a high number of copies in the area of differentiation.  We adopted a similar approach as for the assembly of the thick-billed murre genome (Tigano *et al.* 2018).  We combined a short-insert shotgun library (320 bp) with three mate-pair libraries with inserts of 3, 5 and 10 kb, respectively.  All libraries were prepared and sequenced at Genome Quebec Innovation Center (McGill University, Montreal). The shotgun library was generated robotically using the KAPA HTP Library Preparation Kit Illumina® platforms (Kapa Biosystems) following the manufacturer's recommendations. Fragments 320 bp long were selected robotically using SPRIselect beads (Beckman Coulter, Brea, CA). The three mate-pair libraries were generated using the Nextera Mate Pair Sample Prep Kit (Illumina Inc., San Diego, CA) as per the manufacturer's recommendations. Size selection of fragments was performed by excising 3, 5 and 10 kb bands from 0.5% agarose gels.  All libraries were sequenced using two lanes of Illumina HiSeq2500 V4 paired end 125bp. One lane was reserved for the shotgun library while the mate-pair libraries were pooled and sequenced in the second lane.

We removed Illumina adapters from raw reads using Trimmomatic v. 0.36 (Bolger et al. 2014) and proceeded with *de novo* genome assembly using Platanus (Kajitani *et al.* 2014), a genome assembler developed for highly heterozygous and repetitive genomes. We assembled the short-insert reads in contigs, scaffolded the resulting preliminary assembly with long-insert mate-pair reads, and used all data to close gaps using default settings using the *assemble*, *scaffold* and *gap_close* modules in Platanus, respectively. We retained only scaffolds longer than 500 bp and calculated assembly summary statistics (Table S2). After the first run, major breaks in the assembly were evident at the breakpoints of the CNV despite high overall assembly contiguity (N50=12 Mb, Table S2). We then reran Platanus several times with progressively smaller initial k-mers and increasing maximum difference for bubble crush, but nonetheless the area of interest could not be assembled. The final genome assembly lacked the genomic area of interest and could not be used as a reference genome; however, it provided further evidence for the location of the CNV in the common murre genome.

**Table S1 | Average linkage disequilibrium measured with $r^2$ in the 200 kb area surrounding the FST peak.** All comparisons were statistically significant (Wilcoxon signed-rank test: p-value < 0.001).

| $r^2$ | Bridled common murres | Unbridled common murres |
|---|---|---|
| Peak | 0.1036 | 0.4112 |
| Area surrounding the peak | 0.0920 | 0.0791 |

**Table S2 | Assembly statistics for the *de novo* assembly of the genome of an unbridled common murre included in this study.**

| Assembly statistics | Unbridled common murre genome assembly |
|---|---|
| Number of scaffolds | 21,199 |
| Total size of scaffolds | 1,173,179,815 bp |
| Longest scaffold | 39,499,335 bp |
| Mean scaffold size | 55,341 bp |
| N50 scaffold length | 12,011,034 bp |
| L50 scaffold count | 32 |
| GC content | 40.46% |
| Gaps (stretches of Ns) | 4.18% |

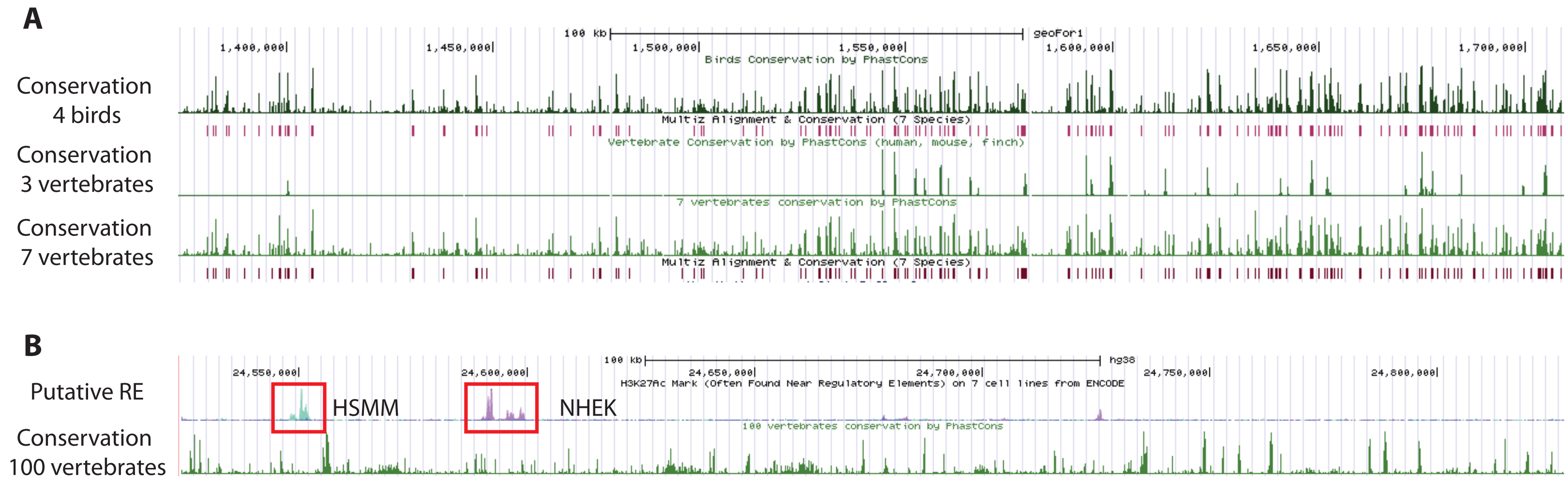
A
Conservation 4 birds
Conservation 3 vertebrates
Conservation 7 vertebrates
100 kb
geoFor1
1,400,000
1,450,000
1,500,000
1,550,000
1,600,000
1,650,000
1,700,000
Birds Conservation by PhastCons
Multiz Alignment & Conservation (7 Species)
Vertebrate Conservation by PhastCons (human, mouse, finch)
7 vertebrates conservation by PhastCons
B
Putative RE
Conservation 100 vertebrates
hg38
24,550,000
24,600,000
24,650,000
24,700,000
24,750,000
24,800,000
H3K27Ac Mark (Often Found Near Regulatory Elements) on 7 cell lines from ENCODE
100 vertebrates conservation by PhastCons
HSMM
NHEK

**Figure S1** Phastcons conservation scores and ENCODE scores from UCSC Genome Browser along the intergenic area. A) Conservation scores (CS) based on 4 bird species (zebra finch, budgerigar, chicken, turkey) and associated conservation elements (CE) in red underneath; CS based on human, mouse and finch; CS based on all 7 species of birds and mammals combined with associated CE. All phastcons analyses are based on multiple alignments to the medium ground finch genome. B) ENCODE scores based on the human genome. High ENCODE scores are generally associated with regulatory elements. ENCODE scores are particularly high in epidermal keratinocytes and skeletal muscle myoblasts (NHEK and HSMM cell lines respectively). CS scores across 100 vertebrate species aligned to the human genome.

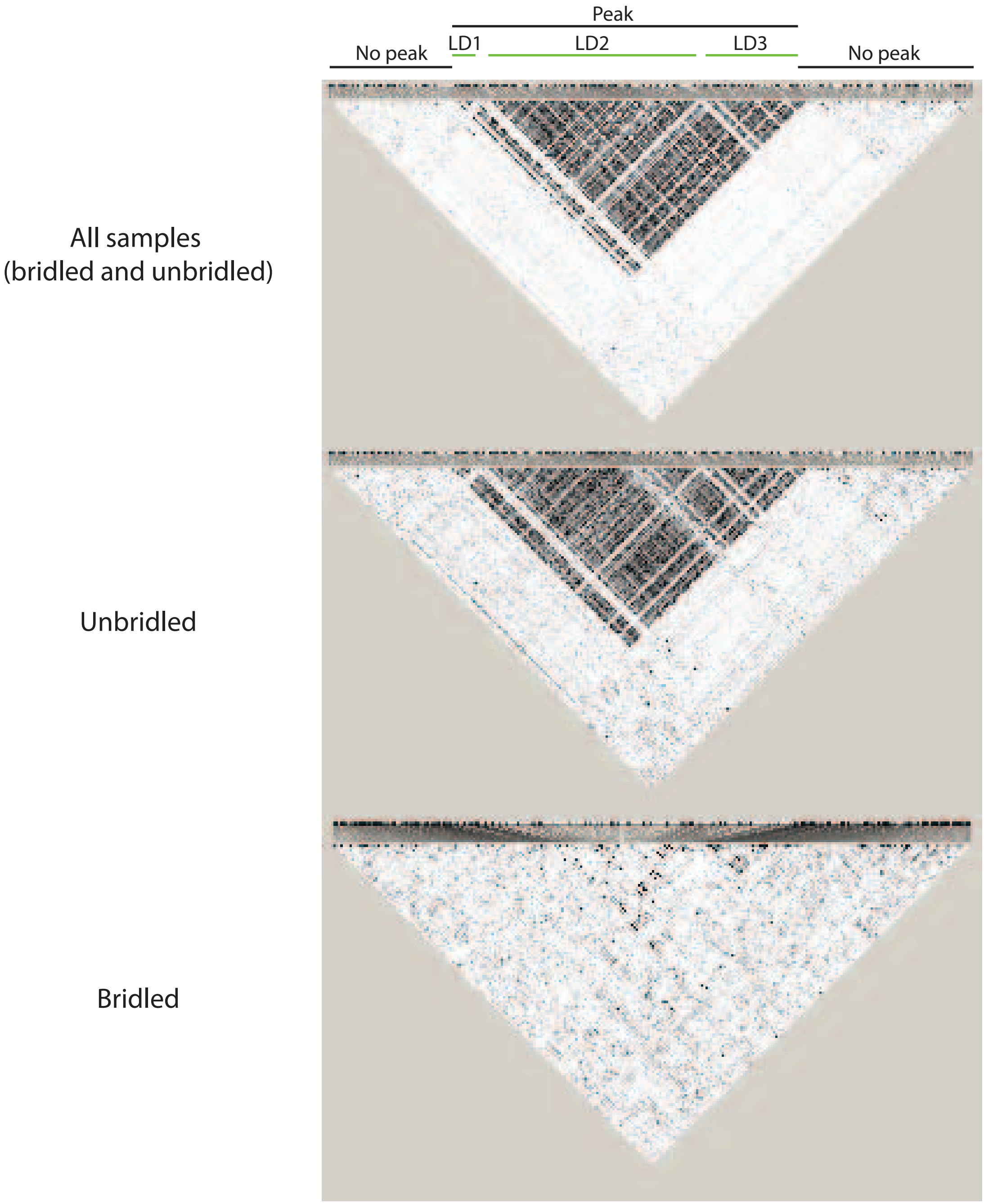


**Figure S2** Pattern of linkage disequilibrium (LD) based on $r^2$ across 200 kb including the area of differentiation. Plots of LD for all samples (n=20), only unbridled (n=10) and only bridled (n=10). The colour intensity (black) indicates the strength of LD in a pairwise comparison between two polymorphic sites.
